# The 5-HT_1A_ receptor agonist NLX-112 rescues motor swimming deficits in Spinocerebellar Ataxia type 3 mice

**DOI:** 10.1101/2025.09.04.674027

**Authors:** D Cunha-Garcia, B Ferreira-Lomba, S Guerreiro, A Vidinha-Mira, D Monteiro-Fernandes, MA Varney, MS Kleven, P Maciel, A Teixeira-Castro, S Duarte-Silva, A Newman-Tancredi

## Abstract

**Introduction:** Spinocerebellar ataxia 3 (SCA3) is a rare neurodegenerative disorder which causes progressive motor disturbances. There is no approved drug treatment but selective activation of serotonin 5-HT_1A_ receptors may be a promising therapeutic strategy to attenuate ataxia symptoms.

**Methods:** NLX-112, a highly selective 5-HT_1A_ full agonist, was tested in the CMVMJD135 transgenic mouse model of SCA3. NLX-112 (1.25 and 5 mg/kg/day) was administered BID intraperitoneally for 14 weeks starting when the mice were 12 weeks of age, i.e., after ataxia signs had become established. The motor swimming test (MST), where mice are required to swim to a raised platform, was used to evaluate the motor behavior of SCA3 mice and their performance was compared with that of wild-type (WT) mice.

**Results:** Both doses of NLX-112 were well tolerated by the SCA3 mice, as assessed by welfare parameters. In the MST, the latency of SCA3 mice to reach the platform was significantly longer than that of WT mice. However, when SCA3 mice were treated with either 1.25 or 5 mg/kg/day of NLX-112, they showed robust improvement of motor performance, with swimming latencies which were similar to those of WT mice. This effect of NLX-112 was maintained throughout the period of the study.

**Conclusions:** The improved motor function of SCA3 mice when treated with NLX-112 supports its investigation as a drug candidate for the treatment of ataxia and related movement disorders.

**Graphical Abstract:** 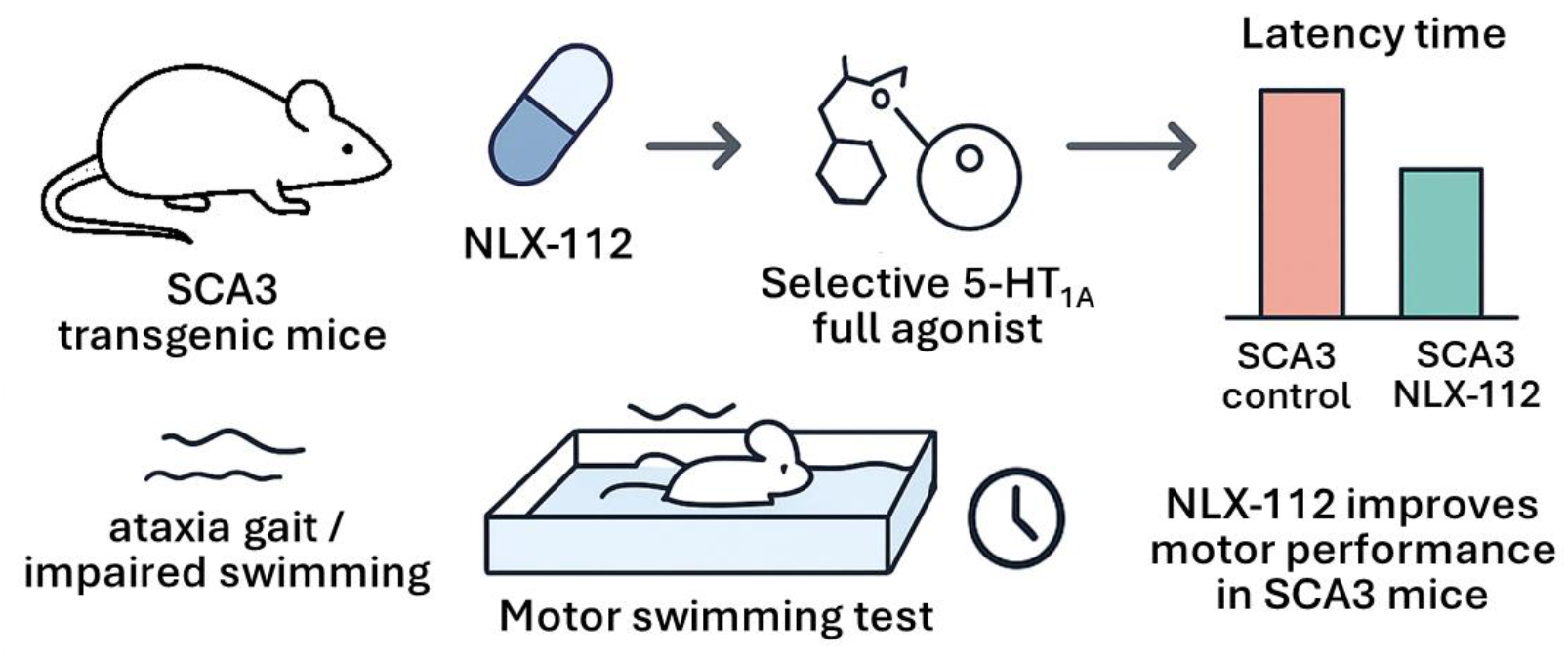

**Highlights:**

- Spinocerebellar ataxia type 3 (SCA3) is marked by progressive motor disturbances
- Targeting the serotonergic system is a promising strategy to treat SCA3
- Transgenic SCA3 mice were treated with NLX-112, a selective 5-HT_1A_ agonist
- Chronic NLX-112 normalized performance of mice in the motor swimming test
- NLX-112 could constitute a drug candidate for treatment of the ataxia disorders

## Introduction

Spinocerebellar ataxia type 3 (SCA3) is the most common autosomal dominant inherited ataxia (Schols et al., 2004). This neurodegenerative disease is caused by an unstable expansion of a cytosine-adenine-guanidine (CAG) trinucleotide within the coding region of the *ATXN3* gene, which is translated into an abnormal ataxin-3 (ATXN3) protein carrying an extended polyglutamine (polyQ) tract. CAG repeat size in healthy individuals ranges from 10 to 44, whereas in SCA3 patients it varies from 61 to 87 and is inversely correlated with the patients’ age at disease onset (Maciel et al., 1995). A core feature of SCA3 is progressive ataxia, characterized by loss of motor coordination, affecting gait, balance, fine movements and speech. Moreover, rigidity, spasticity, dystonia and muscle weakness are often present, as are ophthalmoplegia and nystagmus (Coutinho and Sequeiros, 1981).

The selective serotonin reuptake inhibitor (SSRI) citalopram, which inhibits 5-HT transporter (SERT) function to increase 5-HT synaptic availability, reduced mutant ATXN3 aggregation and improved motor function in *C. elegans* and mouse models of SCA3 (Ashraf et al., 2019; Teixeira-Castro et al., 2015). The beneficial effect in *C. elegans* was dependent on MOD-5, the SERT homologue, and on SER-4, the *C. elegans* homologue of the mammalian 5-HT_1A_ receptor (5-HT_1A_R) (Teixeira-Castro et al., 2015). These observations provide a robust rationale to explore the effects of targeting serotonin neurotransmission in SCA3, and particularly the 5-HT_1A_R, which is one of the major serotonin receptors in the CNS, as targets for treatment of ataxia.

Indeed, 5-HT_1A_Rs have attracted interest as a promising therapeutic target for SCA3. Tandospirone, a 5-HT_1A_R receptor partial agonist, showed encouraging clinical results in SCA3 and SCA6 patients, reducing cerebellar ataxia and other disease-related symptomatology (Takei et al., 2005; Takei et al., 2010). However, tandospirone has only modest receptor specificity (Hamik et al., 1990), which may limit its efficacy and lead to side effects. In contrast, NLX-112 (befiradol, F-13640) is an exceptionally selective agonist with high affinity for 5-HT_1A_Rs (Colpaert et al., 2002; Newman-Tancredi et al., 2017) and shows efficacy in multiple models of movement disorders: it reverses haloperidol-induced dystonia and levodopa-induced dyskinesia (LID) in parkinsonian rats (Iderberg et al., 2015; McCreary et al., 2016) and reduces dyskinesia and parkinsonism in MPTP-treated marmosets and macaques (Depoortere et al., 2020; Fisher et al., 2020). Moreover, a recent proof-of-concept Phase 2A trial of NLX-112 found that it significantly reduced levodopa-induced dyskinesias and parkinsonism in subjects with Parkinson’s disease (Svenningsson et al., 2025), suggesting that it may also be efficacious in alleviating other movement disorders, such as SCA. Indeed, acute and chronic treatment with NLX-112 rescued motor function and suppressed mutant ATXN3 aggregation in a *C. elegans* model of SCA3, supporting further investigation of its effects in animal models of SCA (Pereira-Sousa et al., 2021). The present study therefore explored the effects of NLX-112 in a transgenic mouse model of SCA3, namely the CMVMJD135 mouse strain (Silva-Fernandes et al., 2014a) which closely mimics the gradual appearance and progression of SCA3 symptomatology observed in patients (Duarte-Silva and Maciel, 2018; Esteves et al., 2019; Silva-Fernandes et al., 2014a; Teixeira-Castro et al., 2015). NLX-112 was administered twice daily for 14 weeks, starting at 12 weeks of age, i.e., after the appearance of ataxia signs, which occurs from about 7 weeks of age. We observed that NLX-112 improved motor performance in the SCA3 mice, as assessed by the motor swimming test. These findings support NLX-112’s potential utility as a treatment for movement disorders.

## Methods

### Study design

NLX-112 was administered chronically to the mice via intraperitoneal injections (IP) starting at age 12 weeks, i.e., after the appearance of motor deficits (post-symptomatic treatment initiation), and ended at 27 weeks of age. The animals were randomly allocated to treatment groups at five weeks of age. A total number of 45 animals were used in this study (n=11-12 per experimental group, mixed sexes). All experiments were designed and conducted according to FELASA guidelines and recommendations, considering the 3Rs (Replacement, Reduction and Refinement) and defined humane endpoints. Protocols were approved by the Local Animal Welfare Body of Life and Health Sciences Research Institute (ORBEA-ICVS, reference ORBEA EM/ICVS-I3Bs_007/2019) of the University of Minho, and by the governmental entity *Direção Geral da Alimentação e Veterinária* (DGAV-2022-02-07 003455). The animal facilities and the researchers who worked with mice were licensed by DGAV.

### Mouse Model Generation, Care and Euthanasia

Wild-type (WT, n=11) and CMVMJD135 (i.e., SCA3, n=34) mice littermates (genetic background strain C57BL/6J) of both sexes were used in this study. The experimental groups were defined as follows: WT vehicle (VEH, n=11, 5 males (M)/6 females (F)), SCA3 VEH (n=12, 3M/9F), SCA3 NLX 1.25 mg/kg (n=11, 8M/3F) and SCA3 NLX 5 mg/kg (n=11, 6M/5F). The CMVMJD135 transgenic model expresses the human ATXN3c cDNA variant ubiquitously, regulated by the CMV promoter. A (CAG)_2_CAAAAGCAGCAA(CAG)_129_ repeat tract sequence is carried in the *ATXN3* gene, coding for 135 glutamines (Silva-Fernandes et al., 2014a). DNA extraction, genotyping and CAG repeat determination were performed as previously reported (Silva-Fernandes et al., 2010; Silva-Fernandes et al., 2014a).

The animals, generated in-house, had a Specified Pathogen Free health status and were maintained under standard conditions in the vivarium, namely: artificial 12 h light/dark cycle (lighting from 8 AM to 8 PM), 21 ± 1°C and relative humidity of 50-60 %. A maximum of five mice were housed by genotype and treatment conditions in the Digital Ventilated Cage (DVC®) rack (Tecniplast), which enabled direct monitoring of the mice’s activity in real-time (24 h/7 days), with corncob bedding (Scobis Due, Mucedola S.r.l.), enriched with soft tissues for nesting behavior. The animals were fed with a standardized diet, during gestation, postnatally (4RF25, Mucedola S.r.l.) and after weaning at 3 weeks-old (4RF21, Mucedola S.r.l.). Food and drinking water were available *ad libitum*. Body weight and animal welfare were evaluated weekly. Body temperature of each animal was measured using a thermal rectal probe (Rodent Warmer X1, Stoelting Co.) coated in petroleum jelly at several time points.

### Compound Preparation and Administration

NLX-112 (befiradol fumarate) was provided by Neurolixis SAS. A stock solution was prepared weekly by dissolving in water and then diluted each day in saline prior to IP administration. Each mouse received two injections/day of saline or NLX-112 [0.63 and 2.5 mg/kg *bis in die* (BID); *i*.*e*., 1.25 and 5 mg/kg/day] at 8 AM and at 4 PM. Doses of NLX-112 were chosen based on previous experience (Newman-Tancredi et al., 2022). NLX-112 was administered to SCA3 mice starting at week 12, an age where all the core motor symptoms are fully present (Silva-Fernandes et al., 2014a) (See **Figure 1** for experimental design).

**Figure 1:**
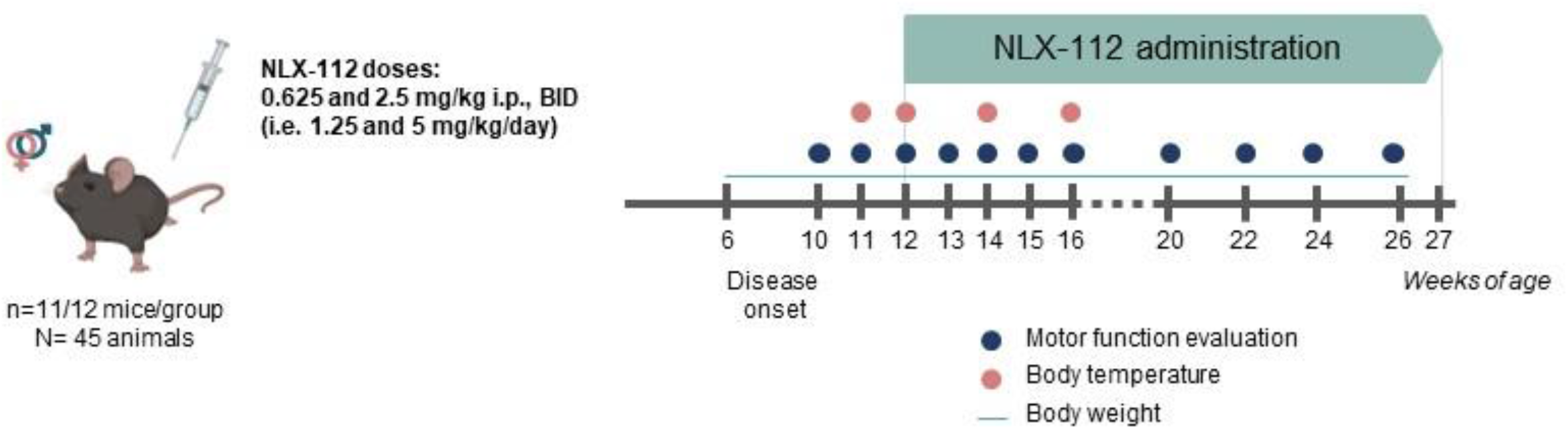
Experimental design for assessment of NLX-112 upon chronic administration to SCA3 mice.

### Behavioral Evaluation

Motor function evaluation was carried out during the light period using the motor swimming test (MST) as described previously (Carter et al., 1999; Silva-Fernandes et al., 2014b). Briefly, the test assesses the time required by the mice to swim from an initial position to a ‘safe’ platform placed at a distance of 60 cm on the surface of the water (water temperature at 23 ± 1 °C). The mice were initially trained on 2 consecutive days (3 trials/animal) and evaluated in the next 3 days (2 trials/animal). The trial was considered invalid if the animals floated or did not swim in a straight line. Testing of the mice began when they were 10 weeks of age and were performed weekly until 20 weeks of age (except at 12 weeks), and then biweekly, until the animals reached 26 weeks of age. The experimenters were blinded to the experimental group.

### Statistical Analysis

Statistical analysis was performed using IBM SPSS Statistics 25 and GraphPad Prism 10.5.0. The sample size applied in each experiment was calculated utilizing G*Power 3.1.9.4, based on power analysis obtained from previous studies, and the effect size was determined based on an improvement of 50 %. The critical value used for significance was *P* ≤ 0.05. The normality of the data was analyzed using descriptive statistics (central tendency, dispersion, kurtosis and skewness) and the Shapiro-Wilk Test (*P* > 0.05) and homogeneity of variances was measured by applying the Levene’s Test (*P* > 0.05). Outliers were removed when the interquartile ranges from the mean deviated more than 1.5. One-Way ANOVA, Two-Way ANOVA and Mixed-design ANOVA were performed to compare group measures, followed by Tukey’s or Sidak post-hoc tests. When the assumption of normal distribution was not met, the results were analyzed by Kruskal-Wallis’s test, followed by Dunn’s post-hoc test.

## Results

Analysis of CAG repeat number found no significant differences between the treatment groups (Supplementary **Figure S1A**), thus excluding a potential confounding effect based on genetic profile of the mice. The mean body temperature of SCA3 mice was significantly lower than that of WT mice but was not modified by either of the doses of NLX-112 (**Figure S1B**).

The mean body weight of SCA3 mice was significantly lower than that of WT mice (**Figure S2**). Treatment with NLX-112 (1.25 mg/kg/day) did not influence body weight of SCA3 mice and did not impact the welfare of the SCA3 mice, as assessed by an absence of adverse effects on animal welfare parameters, including body position, respiratory rate, grooming behavior and hydration.

As concerns the motor swimming test, both WT and SCA3 mice reached the ‘safe’ platform with latencies of about 2s, when tested at ages 10 and 11 weeks, i.e., after ataxia signs had become established but before NLX-112 / vehicle treatment initiation. However, performance of the WT-Vehicle and SCA3-Vehicle groups then diverged: both of these control groups showed decreases in latency, but the WT mice more markedly improved their performance and reached a stable latency of about 1.4 seconds, whereas the SCA3 mice only decreased their latency to 1.6 seconds and then stabilized at about 1.7s (**Figure 2A**), reflecting their ataxic motor impairment.

**Figure 2:**
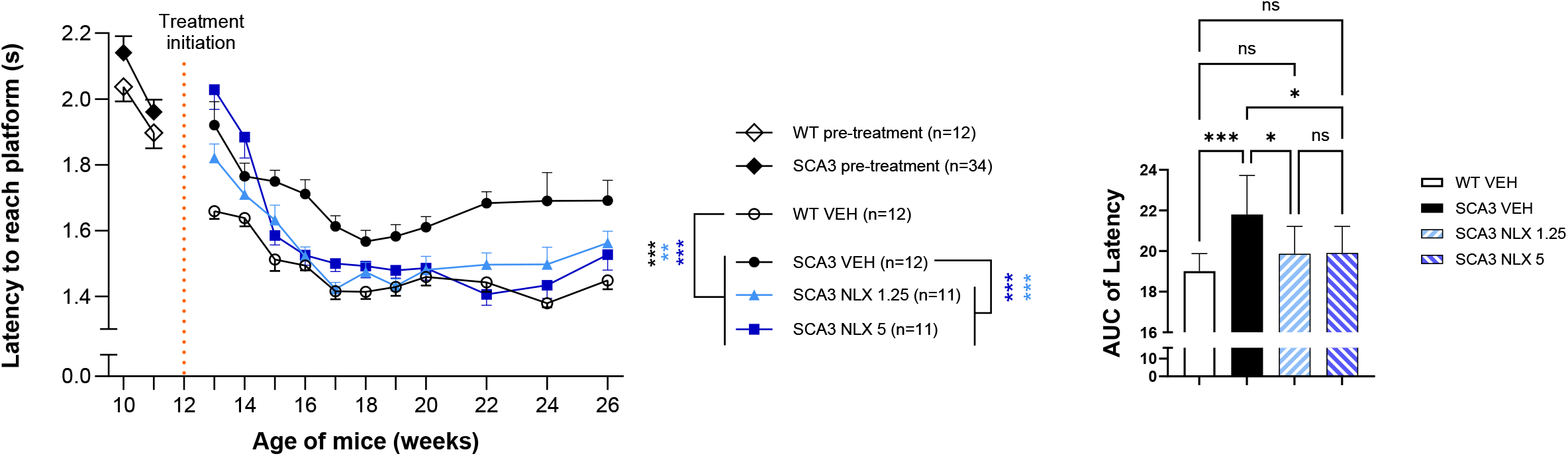
Chronic administration of NLX-112 improved motor function of SCA3 mice as assessed in the motor swimming test. Left panel: Time-course of the latency of the mice to reach a platform placed above the surface of the water. Administration of NLX-112 or vehicle started at week 12 (i.e., after onset of ataxia signs at week 7). Right panel: AUC of total latency times for each group of mice from age 13 to 26 weeks. Time course analysis: two-way ANOVA and mixed-design ANOVA, Tukey, or Sidak post-hoc tests. AUC analysis: one-way ANOVA, Tukey post-hoc test. All data are expressed as group mean ± SEM (* p<0.05, ** p<0.01, *** p<0.001).

SCA3 mice chronically treated with either 1.25 or the 5 mg/kg/day of NLX-112 showed robust improvement in motor performance: their latency to reach the safe platform decreased to levels that, overall, were not significantly different to that of WT mice (see Area Under the Curve data, **Figure 2B**). The improved latency elicited by NLX-112 was maintained throughout the treatment period, although a slight increase was observed at the later time points (weeks 24 and 26), presumably attributable to the worsening motor condition of the mice.

## Discussion

The development of novel treatments for spinocerebellar ataxias constitutes a clear unmet medical need, because there are currently no efficacious approved drugs for this debilitating group of disorders. The present study on NLX-112 (befiradol) suggests that selective targeting of the 5-HT_1A_R could represent a promising strategy for improving symptoms of individuals living with SCA3. The main findings are that NLX-112, administered intraperitoneally BID for 14 weeks starting after the onset of ataxia-related motor deficits, was well tolerated in transgenic male and female SCA3 mice and, importantly, normalized their motor performance, as assessed by the motor swimming test.

As outlined in the Introduction, targeting serotonergic signaling is a promising therapeutic approach for treatment of SCA3 and other forms of ataxia, providing a robust rationale to explore the effects of activating 5-HT_1A_R with a selective and high efficacy agonist. Here, NLX-112 treatment was started when SCA3 mice were aged 12 weeks, i.e., when the ataxia-related motor deficits had become established. This experimental design mimics commonly observed clinical situations, where SCA3 patients are often diagnosed and prescribed treatment after symptoms appear. Chronic treatment with NLX-112 was well tolerated by the SCA3 mice, consistent with the favorable profile of NLX-112 observed in previous chronic studies in rat (McCreary et al., 2016) and in clinical trials. Indeed, NLX-112 has been previously administered to over 600 subjects in various Phase 1 and Phase 2 trials for durations up to 3 months, and was safe and well-tolerated (Neurolixis data on file).

The key observation here is that chronic administration of NLX-112 (1.25 or 5 mg/kg/day), starting after symptom onset, normalized motor performance of SCA3 mice as assessed by the motor swimming test. This test provides an indication of muscle strength and motor performance of the mice (Carter et al., 1999; Silva-Fernandes et al., 2014b), two parameters that are progressively impacted by SCA3. All the mouse treatment groups in the present study (whether WT or SCA3) showed an initial improvement in performance, i.e., the latency to reach the ‘safe’ platform decreased over the first 4 or 5 weeks of treatment (i.e., from when the mice are aged 13 weeks until they are aged to 17 or 18 weeks) reflecting a learning / acquisition phase (Jiang et al., 2017). However, notable differences were observed between the groups: the WT mice substantially improved their performance, whereas the SCA3 mice showed only partial improvement and then deteriorated somewhat as they aged. In contrast, mice treated with either 1.25 or 5 mg/kg/day of NLX-112 showed latency times which were significantly shorter compared to vehicle-treated SCA3 mice and similar to those of WT mice. The improved latency elicited by NLX-112 was maintained throughout the treatment period, even when they reached 22 to 26 weeks of age, when motor deficits become increasingly disabling. These observations are concordant with those found previously in nematode worm experiments, where NLX-112 improved motility in a *C. elegans* model of SCA3, (Pereira-Sousa et al., 2021), in experiments on non-human primates, where NLX-112 significantly reduced motor disability in MPTP-lesioned marmosets (Fisher et al., 2020), and in human subjects with Parkinson’s disease, where NLX-112 significantly reduced UPDRS (Unified Parkinson’s Disease Rating Scale) motor disability scores (Svenningsson et al., 2025).

As concerns the mechanism of action of NLX-112, electrophysiology experiments showed that it strongly activates 5-HT_1A_ autoreceptors in the dorsal raphe nucleus, thereby inhibiting the firing of 5-HT neurons. It also activates 5-HT_1A_ heteroreceptors, thereby increasing dopamine release and increasing the firing of glutamatergic pyramidal neurons in cortical regions (Llado-Pelfort et al., 2012). Moreover, brain imaging experiments using ^18^F-labeled NLX-112 found labelling in basal ganglia and cerebellum (Colom et al., 2020; Vidal et al., 2018), areas which are strongly implicated in the neuropathology of SCA3 and other movement disorders (Klockgether et al., 2019; Sival et al., 2022). It may therefore be surmised that NLX-112 exerts its beneficial effects by targeting several brain regions involved in motor performance, although this warrants further investigations to determine their precise roles.

The present data raise important questions concerning the broader effects of targeting 5-HT_1A_ receptors in SCA3. Firstly, the study investigated the capacity of NLX-112 to attenuate motor deficits after they were already established in the SCA3 mice. It would be informative to determine whether NLX-112 could also prevent the development of motor symptoms, by starting treatment of the mice before onset of SCA-related deficits. Secondly, although the motor swimming test is useful for assessing motor performance in SCA mice, it would be desirable to test whether NLX-112 treatment also has a beneficial effect on other behavioral models, notably the beam walking test, which is a pertinent for detecting ataxia-related motor coordination deficits. Thirdly, the present study used twice-daily intraperitoneal (i.e., pulsatile) administration as a drug delivery route, implying that there were fluctuations of NLX-112 exposure in blood and brain of the SCA3 mice (the half-life of NLX-112 in human is >24 hours but about 2-4 hours in rodents; data on file). It would therefore be interesting to explore the effects of the drug using a delivery route which provides a more stable exposure to NLX-112, such as in drinking water. It is known that NLX-112 is chemically stable for several weeks in solution, as shown in previous studies where it was administered subcutaneously by osmotic minipumps (Bruins Slot et al., 2003). Finally, 5-HT_1A_ receptor agonists are known to exert neuroprotective properties in models of movement disorders (Miyazaki and Asanuma, 2017). It would be interesting to determine whether NLX-112 protects against neuronal damage in SCA3 mice, particularly as concerns dopaminergic neurons in the Substantia nigra. It should be noted that the authors have carried out an extensive investigation of the above questions and the results are disclosed as a pre-print (Ferreira-Lomba, 2025).

Overall, the present results support the assertion that selective activation of 5-HT_1A_R could be a useful therapeutic approach for SCA3 and other movement disorders. As mentioned above, NLX-112 exhibits antiparkinsonian and antidyskinetic properties in animal models (Depoortere et al., 2020; Fisher et al., 2020; Iderberg et al., 2015; McCreary et al., 2016) and significantly reduced both dyskinesia and parkinsonism in a proof-of-concept clinical trial (Svenningsson et al., 2025), suggesting that it has broad activity to relieve motor symptoms in diverse movement disorders. In this context, NLX-112 has been granted Orphan Drug designation as a treatment for Spinocerebellar Ataxia in both the European Union and in the United States (Designation numbers: EU/3/24/2951 and DRU-2025-10752), supporting its clinical development.

Taken together, these data on SCA3 mice suggest that selective, pulsative, activation of 5-HT_1A_R by NLX-112, starting at a post-symptomatic stage of the disease, can improve its motor symptoms in a sustained manner over an extended period. The study supports further investigation of NLX-112 as a treatment for spinocerebellar ataxias and related movement disorders.

## Acknowledgments

We are grateful to all the members and some ex-members of the Translational Neurogenetics team for the critical analysis of the data and helpful discussions.

## Authors’ Roles

Daniela Cunha-Garcia: execution, analysis, writing

Bruna Ferreira-Lomba: execution, writing

Sara Guerreiro: execution, writing

André Vidinha-Mira: execution Daniela Monteiro-Fernandes: execution

Mark A Varney: design, funding acquisition

Mark S Kleven: design, management, supervision, editing of final version of the manuscript

Patrícia Maciel: design, management, supervision, funding acquisition, editing of final

Andreia Teixeira-Castro: execution, design, management, supervision, editing of final version of the manuscript.

Sara Duarte-Silva: design, execution, analysis, writing, management, supervision, editing of final version of the manuscript.

Adrian Newman-Tancredi: design, funding acquisition, management, supervision, writing, editing of final version of the manuscript.

## Data Sharing

Data will be made available upon reasonable request.

