## Supplemental Figures for "The 5-HT_1A_ receptor agonist NLX-112 rescues motor swimming deficits in Spinocerebellar Ataxia type 3 mice"

**Figure S1 (Supplemental Information):** Comparison of CAG repeats and body temperature between groups. Left panel: The CAG repeat length did not significantly differ between treatment groups. Right panel: SCA3 mice exhibited significantly lower body temperature than WT mice. However, chronic administration of NLX-112 to SCA3 mice, assessed over 4 weeks (i.e., from age 12 to 16 weeks) did not modify their core body temperature.

CAG repeat analysis by Kruskal-Wallis test, Dunn's post-hoc test. Body temperature analysis by Two-way ANOVA or mixed-design ANOVA, Tukey, or Sidak post-hoc tests. All data are expressed as group mean ± SEM (* p<0.05, ** p<0.01, *** p<0.001).

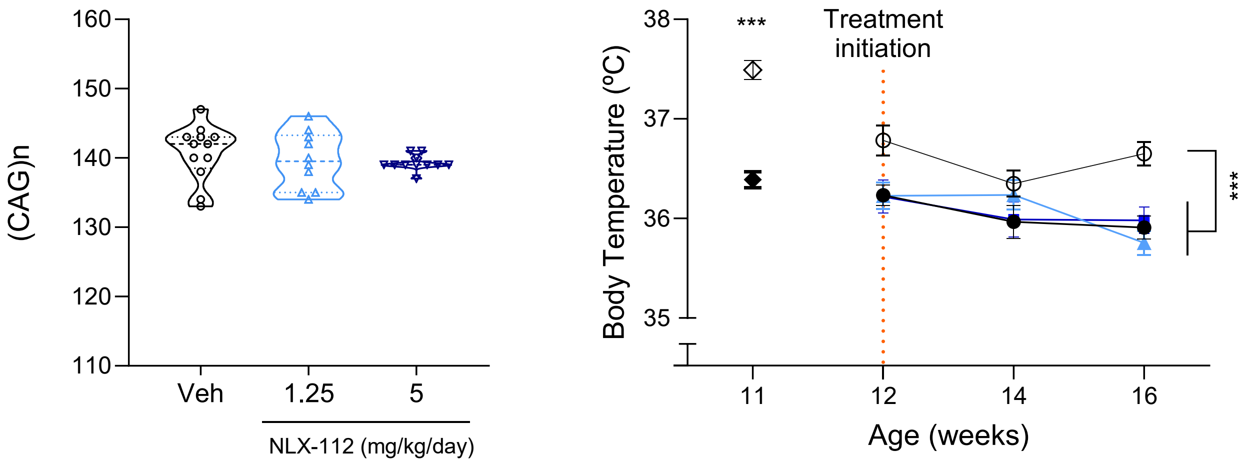

**Figure S2 (Supplemental Information):** Changes in body weight of WT and SCA3 mice. NLX-112 (1.25 or 5 mg/kg/day) did not have observable effects on body weight of SCA3 mice compared to vehicle-treated SCA3 controls (n=11-12 per group).

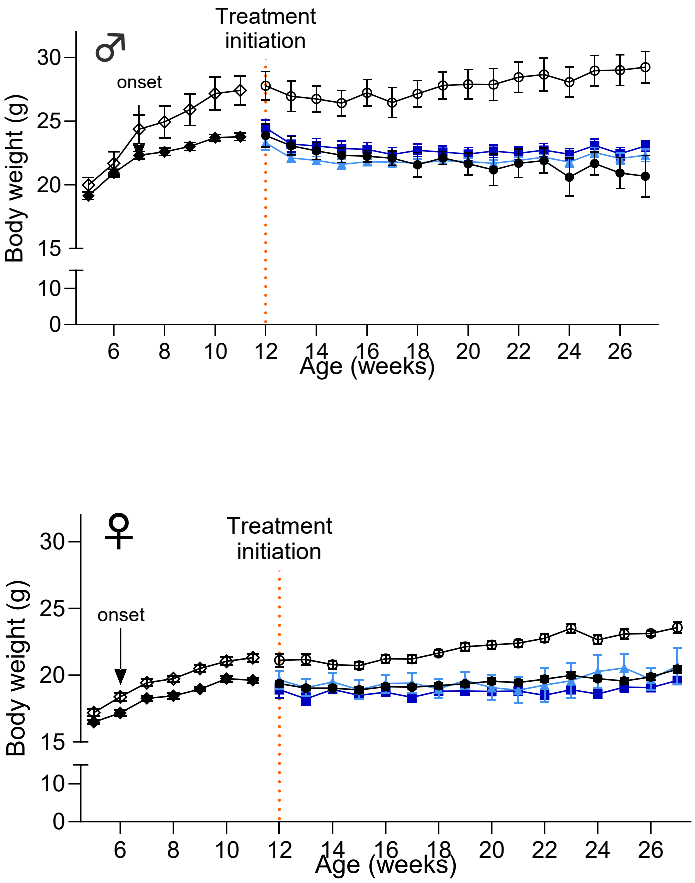
